# Undermining the cry for help: The phytopathogenic fungus *Verticillium dahliae* secretes an antimicrobial effector protein to undermine host recruitment of antagonistic *Pseudomonas* bacteria

**DOI:** 10.1101/2025.06.09.658588

**Authors:** Anton Kraege, Wilko Punt, Andrea Doddi, Jinyi Zhu, Natalie Schmitz, Nick C. Snelders, Bart P.H.J. Thomma

## Abstract

During pathogen attack, plants recruit beneficial microbes in a “cry for help” to mitigate disease development. Simultaneously, pathogens secrete effectors to promote host colonization through various mechanisms, including targeted host microbiota manipulation. Here, we characterize the Av2 effector of the vascular wilt fungus *Verticillium dahliae* as a suppressor of the cry for help. Inspired by *in silico* antimicrobial activity prediction, we investigated the antimicrobial activity of Av2 *in vitro*. Furthermore, its role in *V. dahliae* virulence was assessed through microbiota sequencing of inoculated plants, microbial co-cultivation assays, and inoculations in a gnotobiotic plant cultivation system. We show that Av2 inhibits bacterial growth, and acts as a virulence factor during host colonization. Microbiota sequencing revealed involvement of Av2 in suppression of *Pseudomonas* spp. recruitment upon plant inoculation with *V. dahliae*, suggesting that Av2 suppresses the cry for help. We show that several *Pseudomonas* spp. are antagonistic to *V. dahliae* and sensitive to Av2 treatment. We conclude that *V. dahliae* secretes Av2 to supress the cry for help by inhibiting the recruitment of antagonistic *Pseudomonas* spp. to pave the way for successful plant invasion.

## INTRODUCTION

Plants associate with a plethora of microbes above and below ground, collectively called the microbiota, that can positively impact plant productivity and health (Berendsen et al., 2018; Trivedi et al., 2020). Through the secretion of root exudates plants shape their microbiota and actively recruit beneficial microbes to mitigate biotic and abiotic stresses (Berendsen et al., 2018; López et al., 2008). In the face of pathogen attack, plants can modify these exudates to selectively attract protective microbes in order to limit disease progression. This targeted recruitment in response to pathogen infection is known as the plant’s “cry for help” (Berendsen et al., 2018; Liu et al., 2024; Spooren et al., 2024; Yuan et al., 2018). For example, cucumber plants increase the exudation of tryptophan during *Fusarium oxysporum* infection, which promotes the recruitment of beneficial *Bacillus amyloliquefaciens* that can mitigate disease progression(Liu et al., 2017).

Ultimately, the cry for help recruitment of beneficial microbes may have a legacy effect in cases when it leads to an increased population of these microbes in the soil, resulting in the establishment of disease-suppressive soils that protect future plants grown in the same soil (Mesny et al., 2024; Rolfe et al., 2019). However, the development of such disease suppressiveness typically requires years and many plant generations to fully establish (Rolfe et al., 2019). Arguably, the most famous example of such legacy effect concerns the decline of the take-all disease in wheat over years of monoculture that has been associated with the recruitment of 2,4-diacetylphloroglucinol-producing *Pseudomonas* spp. (Raaijmakers & Weller, 2007).

To detect pathogens, plants have evolved a complex immune system that recognises a multitude of microbe-derived molecules to activate appropriate defence responses (Jones & Dangl, 2006). Initial immune responses are triggered upon recognition of conserved microbe-associated molecular patterns (MAMPs), such as chitin or flagellin, by plant membrane-localised pattern recognition receptors that activate pattern-triggered immune (PTI) responses (Cook et al., 2015; Jones & Dangl, 2006). In response, host-adapted pathogens have evolved strategies to suppress or overcome such PTI responses, which includes the secretion of virulence factors, also known as effectors (Rovenich et al., 2014). In turn, particular host genotypes evolved to recognize effectors, or their activities, by resistance (R) proteins that include cell surface and cytoplasmic receptors that activate effector-triggered immunity (ETI) (Cook et al., 2015; Jones & Dangl, 2006).

Most effectors characterized to date deregulate host immune responses or target other aspects of host physiology through various biochemical activities and mechanisms (Rovenich et al., 2014). For example, the effector Ecp6 is secreted by *Cladosporium fulvum* to sequester chitin oligosaccharides that are released from its cell walls to prevent recognition by chitin immune receptors (Sánchez-Vallet et al., 2013). Intriguingly, several research groups have recently uncovered a novel function of effectors besides the modulation of host physiology, by showing that several pathogens secrete effectors that target host-associated microbiota through the display of selective antimicrobial activity in order to promote host colonisation (Gómez-Pérez et al., 2023; Snelders et al., 2020).

Several antimicrobial effectors have been functionally characterized in the soil-borne fungus *Verticillium dahliae*, a presumed asexual filamentous fungus that causes vascular wilt disease on hundreds of host plants (Fradin & Thomma, 2006). The fungus generates genetic diversity through largescale chromosomal rearrangements and segmental duplications, leading to hypervariable regions between *V. dahliae* strains that are called adaptive genomic regions (AGRs) (Cook et al., 2020; de Jonge et al., 2013; Faino et al., 2016). These AGRs are enriched in repeats and in effector genes, and display a unique chromatin profile that sets these regions apart from core genomic regions (D. E. Cook et al., 2020). Interestingly, despite being dispersed across the genome, these AGRs were found to physically interact in the nucleus, possibly contributing to their differential behaviour (Torres et al., 2024). Overall, similar to other filamentous pathogens, *V. dahliae* has a compartmentalised genome containing AGRs with increased plasticity when compared with core genomic regions, an observation often referred to as a “two-speed genome” (Raffaele & Kamoun, 2012; Torres et al., 2021).

The first *V. dahliae* effector for which antimicrobial activity was shown is the AGR-encoded lineage-specific effector Ave1 that was identified by comparative genomics between *V. dahliae* strains that are controlled by *Ve1-*mediated resistance in tomato and resistance-breaking strains (de Jonge et al., 2012). Besides being recognized by the tomato Ve1 immune receptor as avirulence factor, Ave1 contributes to fungal virulence on plants lacking Ve1 by targeting antagonistic Sphingomonadales (Snelders et al., 2020). Notably, Ave1 is not the only *V. dahliae* effector protein with antibacterial activity, as a search for effectors with homology to known antimicrobial proteins within the *V. dahliae* secretome yielded the AMP2 effector that is expressed in soil extract. AMP2 revealed complementary activity to Ave1, suggesting that *V. dahliae* exploits different effectors to cope with the diversity of microbial competitors in soil (Snelders et al., 2020). The antimicrobial activity of *V. dahliae* effector proteins is not restricted to bacteria, as the defensin-like effector AMP3 was found to target the mycobiome (Snelders et al., 2021). Intriguingly, and in contrast to Ave1 and AMP2, AMP3 is exclusively expressed at late infection stages when resting structures are formed in decaying plant tissue while host immune responses fade and opportunists and fungal decay organisms invade host tissues (Snelders et al., 2021).

Over the years only two *R* loci were identified that confer resistance against *V. dahliae* in tomato. Besides the recognition of Ave1 by the Ve1 receptor, the fungal effector Av2 is recognised in *V2* tomato plants, although the corresponding *R* gene has not yet been cloned (Chavarro-Carrero et al., 2021; Usami et al., 2017). Similar to Ave1, Av2 is a small (73 amino acid mature protein; net charge +1.8) secreted protein, produced only by a subset of *V. dahliae* strains. Apart from homologues found in other *Verticillium* spp., the only homologues of this effector where found in the *Fusarium* genus (Chavarro-Carrero et al., 2021). *V. dahliae Av2* occurs in two allelic variants that differ in one non-synonymous single nucleotide polymorphism (SNP) that are both recognised in *V2* plants, and so far its intrinsic function for the pathogen has remained enigmatic (Chavarro-Carrero et al., 2021).

## MATERIALS AND METHODS

### Detection of *V. dahliae Av2* expression in soil extract

For each treatment, 10^6^ conidiospores of *Verticillium dahliae* strain JR2 were added to 10 mL potato dextrose broth (PDB) and incubated while shaking with 130 rpm at 28°C for 2 days. Subsequently, the mycelium was collected using sterilized miracloth (Merck, Darmstadt, Germany) and washed with sterilized water. Next, the mycelium was transferred to new flasks containing 10 mL of soil extract that was prepared by adding 40 g of potting soil to 200 mL of sterilized water followed by incubation at room temperature for 2 days, after which soil particles were pelleted by centrifugation for 30 minutes at 4,000 x g and the supernatant was collected. The flasks were then incubated while shaking with 130 rpm at 28°C for 5 days. Next, mycelium was recollected using sterilized miracloth and washed with sterilized water. RNA was extracted using TRIzol reagent (Thermo Fisher Scientific, Waltham, Massachusetts, USA) of which 1 µM was transcribed into cDNA using the PrimeScript™ RT reagent Kit with gDNA Eraser (Takara Bio USA, San Jose, CA, USA). Real-time PCR was performed using SsoAdvance Universal SYBR Green Supermix (BioRad, Hercules, California, USA) and the expression of effector genes was normalized using the *V. dahliae* glyceraldehyde 3-phosphate dehydrogenase gene (*VdGAPDH*) as a reference.

### Heterologous expression of Av2 homologs

The *Av2* alleles encoding *V. dahliae* Av2 and Av2^V73E^ (from strains TO22 and JR2, respectively) and their homologs from *Fusarium oxysporum* f.sp. *pisi* and *F. redolens* (*Fop*Av2 and *Fr*Av2, respectively), were codon-optimized for expression in *E. coli* and cloned into the pET15b vector (Merck, Darmstadt, Germany) such that the proteins are produced without a signal peptide and as fusion protein with an N-terminal His_6_ tag. All vectors were ordered from BioCat GmbH (Heidelberg, Germany). While *VdAv2* and Av2^V73E^ were produced in *E. coli* strain BL21 (Thermo Fisher Scientific, Waltham, Massachusetts, USA), *Fop*Av2 and *Fr*Av2 were produced in *E. coli* strain SHUFFLE T7 (New England Biolabs, Ipswich, Massachusetts, USA). A pre-inoculum of bacterial cultures was incubated overnight in Luria and Bertani (LB) broth supplemented with 50 µg/mL ampicillin at 37°C for BL21 and at 30°C for SHUFFLE T7 while shaking at 170 rpm. Subsequently, the pre-inoculum was transferred to 1 L of LB supplemented with ampicillin (50 µg/mL) and incubated at 37°C (BL21) or 30°C (SHUFFLE T7) until the optical density at 600 nm (OD_600_) reached 0.6-0.8. Next, isopropyl-1-thio-β-D-galactopyranoside (IPTG) was added to a final concentration of 1 mM and the culture was incubated for 4 h at 37°C (BL21) or 30°C (SHUFFLE T7). Next, cells were pelleted through centrifugation (21,000 x g) at 4°C and resuspended in 6 M guanidine, 10 mM TRIS-HCl and 10 mM β-mercaptoethanol (pH 8.0) and incubated overnight at 4°C while rotating continuously. After centrifugation at 21,000 x g for 30 minutes, proteins were purified from the supernatant by immobilized metal affinity chromatography (IMAC) on a custom packed 5 mL Ni^2+^ CYTIVIA column (volume, supplier) with His60 Ni Superflow Resin (Takara Bio USA, San Jose, CA, USA). Fractions containing the protein of interest were identified by sodium dodecyl sulphate-polyacrylamide gel electrophoresis (SDS-PAGE) analysis, combined and dialysed in a stepwise fashion. To this end, in the protein was first dialyzed for 18 h against 4 M guanidine, 50 mM BIS-TRIS, 10 mM reduced glutathione, 2 mM oxidized glutathione (pH 7.0), and subsequently in 18 h steps against 3 M guanidine, 50 mM BIS-TRIS, 10 mM reduced glutathione, 2 mM oxidized glutathione (pH 6.5), followed by 2 M guanidine, 100 mM BIS-TRIS, 250 mM ammonium sulfate, 10 mM reduced glutathione, 2 mM oxidized glutathione (pH 6.5). and then by 1 M guanidine, 100 mM BIS-TRIS, 125 mM ammonium sulfate, 10 mM reduced glutathione, 2 mM oxidized glutathione (pH 5.8), followed by 100 mM BIS-TRIS, 125 mM ammonium sulfate, 10 mM reduced glutathione, 2 mM oxidized glutathione (pH 5.8). Ultimately the protein was dialysed against potassium phosphate buffer (pH 6.5). Final protein concentrations were determined with Nanodrop (Thermo Fisher Scientific, Waltham, Massachusetts, USA) based on absorbance at 280 nm and adjusted to a final concentration of 16 μM.

### *In vitro* antimicrobial activity assays

All bacteria used in the assays originated from a tomato culture collection (Punt et al., 2025).After growth of bacterial isolates on tryptone soy agar (TSA) at 28°C, single colonies were selected and grown overnight in tryptone soy low salt broth (TSB LS) at 28°C while shaking at 200 rpm. Subsequently, OD_600_ was adjusted to 0.05 by dilution with TSB LS. Hundred μL of bacterial culture were mixed with 100 μL of Av2 protein variants (16 μM) in clear 96-well plates (BRAND SCIENTIFIC GMBH, Wertheim, Gemany) with three replicates for each treatment. The plates were incubated in a CLARIOstar® plate reader (BMG LABTECH, Ortenberg, Germany) at 28°C with double orbital shaking every 15 minutes (10 seconds at 300 rpm) after which the optical density was measured at 600 nm (Snelders et al., 2020).

### Plant disease assays

Inoculation of tomato plants to determine the virulence of the *V. dahliae* was performed as described previously (Fradin et al., 2009). Briefly, conidiospores of *V. dahliae* wild-type or *Av2* deletion strain were harvested after ten days of cultivation on potato dextrose agar (PDA). The conidiospore suspensions were centrifuged at 10,000 x g for 10 min and the pellets were resuspended in water. This washing step was repeated twice before spores were counted and the concentration was adjusted to 10^6^ conidiospores/mL. For the inoculation, ten-day-old tomato (*Solanum lycopersicum* L.) MoneyMaker plants were carefully uprooted, roots were rinsed in water, and placed into the inoculum for 6 minutes. Next, plants were placed back into soil, and placed in the greenhouse at 22°C during 16-h/8-h day/night periods with maximum 80% relative humidity, and symptoms were monitored at 14 days post inoculation. To this end canopy areas were measured and fungal biomass inside the tomato stem was determined. For the latter, samples were frozen in liquid nitrogen, ground to a fine powder, and DNA was isolated using phenol-chloroform extraction (Chavarro-Carrero et al., 2021). *V. dahliae* biomass was quantified through real-time PCR using *V. dahliae*-specific primers targeting the ITS region of the ribosomal DNA. The tomato rubisco gene was used for sample calibration. Real-time PCRs were conducted using a CFX Opus Real-Time PCR Systems (BioRad, Hercules, California, USA) and the using SsoAdvance Universal SYBR Green Supermix (BioRad, Hercules, California, USA).

### Tomato stem microbiota sequencing

Tomato stems samples were collected, washed with sterile water, frozen in liquid nitrogen and manually ground using mortar and pestle. Total DNA was extracted following a phenol-chloroform based extraction procedure (Chavarro-Carrero et al., 2021) and DNA concentrations were determined using a Qubit fluorometer (Thermo Fisher Scientific, Waltham, Massachusetts, USA). Sequence libraries were prepared following amplification of the V5-V7 region of the bacterial 16S rDNA (799F and 1139R) as described previously (Wippel et al., 2021) and sequenced (paired-end 300 bp) on an Illumina MiSeq V3 Platform (Cologne Center for Genomics, Cologne, Germany). Sample barcoding was done as described previously (Fadrosh et al., 2014).

### Microbiota analysis

Sequencing data were processed using R v.4.2.0. as described previously (Callahan et al., 2016; Snelders et al., 2020). Briefly, reads were demultiplexed using cutadapt (v4.1; Martin, 2011). Afterwards reads were trimmed and filtered to an average paired read length of 412 bp with the Phred score of 30. From the trimmed reads OTUs were inferred using the DADA2 method (v 1.24; Callahan et al., 2016). Taxonomy was assigned using the Ribosomal Database Project (RDP,v 18; Cole et al., 2014). The pyloseq package (v1.40.0; McMurdie & Holmes, 2013) was used to calculate β-diversity (weighted unifrac distance) after the data was normalised with the metagenomeseq package (v1.38.0; Paulson et al., 2013) using cumulative sum scaling. PERMANOVA was performed with the vegan (v2.6-4; Oksanen et al., 2004) package. Differential abundance analysis was done using the DESeq2 package (v1.36.0; Love et al., 2014) using a negative binomial Wald test and a significance P adjusted threshold < 0.05.

### *In vitro* competition assay

Conidiospores of *V. dahliae* strain TO22 and the *VdAv2* deletion strain were harvested from a PDA plate using sterile water and diluted to a concentration of 2 x 10^6^ conidiospores/mL in half-strength Murashige and Skoog (MS) medium (Duchefa, Haarlem, The Netherlands). Plant associated *Pseudomonas* spp. were cultured overnight in half-strength MS medium. Next, overnight cultures were adjusted to OD_600_ of 0.05 in half-strength MS and added to the conidiospores, and 500 μL of the microbial mixture was added into 12-well flat bottom polystyrene tissue culture plate (Sarstedt, Nümbrecht, Germany). Following 48 h of incubation at RT with shaking at 150 rpm, genomic DNA was extracted using the SmartExtract DNA kit (Eurogentec, Maastricht, The Netherlands), and *V. dahliae* biomass was quantified through real-time PCR using *V. dahliae*-specific primers targeting the internal transcribed spacer (ITS) region of the ribosomal DNA (Snelders et al., 2020). A spike-in DNA sequence was added during DNA extraction for sample calibration (Guo et al., 2020). Genomic sequences of the tested *Pseudomonas* spp. (Punt et al., 2025) were used to infer rooted species trees based on single-copy orthologous genes (Emms & Kelly, 2019).

### Gnotobiotic tomato cultivation for *V. dahliae* inoculations

For tomato cultivation, a previously developed Flowpot-system was used (Kremer et al., 2021; Punt et al., 2025). A 1:1 blend of potting soil (Balster Einheitserdewerk, Fröndenberg, Germany) and vermiculite (LIMERA Gartenbauservice, Geldern-Walbeck, Germany) were autoclaved three times on a liquid cycle and filled into 50 mL syringes (Terumo Europe, Leuven, Belgium). To check for substrate sterility, 500 mg of substrate was added to 10 mL of 100 mM MgCl_2_ and shaken at 300 rpm at room temperature for 1 hour. Subsequently, the samples were diluted 1,000-fold and 50 μL of the dilution was plated onto Reasoner’s 2A agar (R2A), TSA and LB agar (LBA), and incubated in darkness at room temperature for 4 days before microbial growth was assessed. The substrate-filled syringes were flushed with 40 mL of sterile H_2_O followed by 40 ml of half-strength MS. Next, surface-sterilized tomato seeds were placed into each syringe and 6 syringes were placed into an autoclaved Microbox container (SacO2, Deinze, Belgium) and placed in the greenhouse at 22°C during 16-h/8-h day/night periods with a maximum of 80% relative humidity. After two weeks, tomato plants were carefully uprooted under sterile conditions and inoculated with 10^6^ conidiospores/mL of either wild-type TO22 or the corresponding *Av2* deletion strain. Subsequently, the plants were placed back into the syringe and the syringes into the container in the greenhouse. Symptoms of disease were scored at 14 days post inoculation. For biomass quantification, stems of the plants were frozen in liquid nitrogen and ground to a fine powder. DNA was isolated using phenol-chloroform extraction (Chavarro-Carrero et al., 2021). *V. dahliae* biomass was quantified through real-time PCR using *V. dahliae* specific primers targeting the ITS region of the ribosomal DNA. The tomato rubisco gene was used for sample calibration. Real-time PCRs were conducted using a CFX Opus Real-Time PCR Systems (BioRad, Hercules, California, USA) and the using SsoAdvance Universal SYBR Green Supermix (BioRad, Hercules, California, USA).

## RESULTS

### Av2 selectively inhibits bacterial growth *in vitro*

Most functionally characterised effectors target host physiology and are strictly *in planta* expressed, while microbiota-manipulating effectors can be expressed *in planta* as well as during fungal life cycle stages outside the plant host (Snelders et al., 2020, 2021). In order to functionally characterize *Av2*, its expression was analysed by querying previously generated RNA sequencing datasets (Cook et al., 2020; de Jonge et al., 2012), revealing that *Av2* is not only expressed during host colonisation (1,695 TPM, 16 dpi, de Jonge et al., 2012) but also during *in vitro* growth on potato dextrose agar (3,256 TPM, 4 day old, Cook et al., 2020). Furthermore, *Av2* is expressed in conditions mimicking soil colonisation (Figure S1a). A similarly broad expression pattern, including expression in soil, has previously been observed for the *V. dahliae Ave1* effector gene (Snelders et al., 2020), suggesting that Av2 may act as an antimicrobial too. Interestingly, *in silico* analysis using the Antimicrobial Peptide Scanner (vr.2; (Veltri et al., 2018)) predicted antimicrobial activity for Av2 with a probability of 99.6%.

In order to validate the predicted antimicrobial activity of Av2 *in vitro*, the two previously identified variants, Av2 and Av2^V73E^, were expressed heterologously in *E. coli*, purified, and used in antimicrobial activity assays. Additionally, Av2 homologues from two *Fusarium* spp. were produced, purified and tested for antimicrobial activity as well. To this end, a panel of ten phylogenetically diverse plant-associated bacteria was incubated with either of the Av2 variants at a concentration of 8 µM, or buffer as a control, and bacterial growth was assessed. Interestingly, three out of ten bacteria showed reduced growth when incubated with either of the two *V. dahliae* Av2 variants, or the *Fusarium* homologues, namely *Bacillus drentensis*, *Pseudoxanthomonas suwonensis* and *Devosia riboflavina* (Figure 1). Importantly, no differences in inhibitory activity were observed between any of the Av2 variants, suggesting they have overlapping activity spectra. Thus, all Av2 proteins display selective antimicrobial activity against bacteria *in vitro*.

**Fig. 1.**
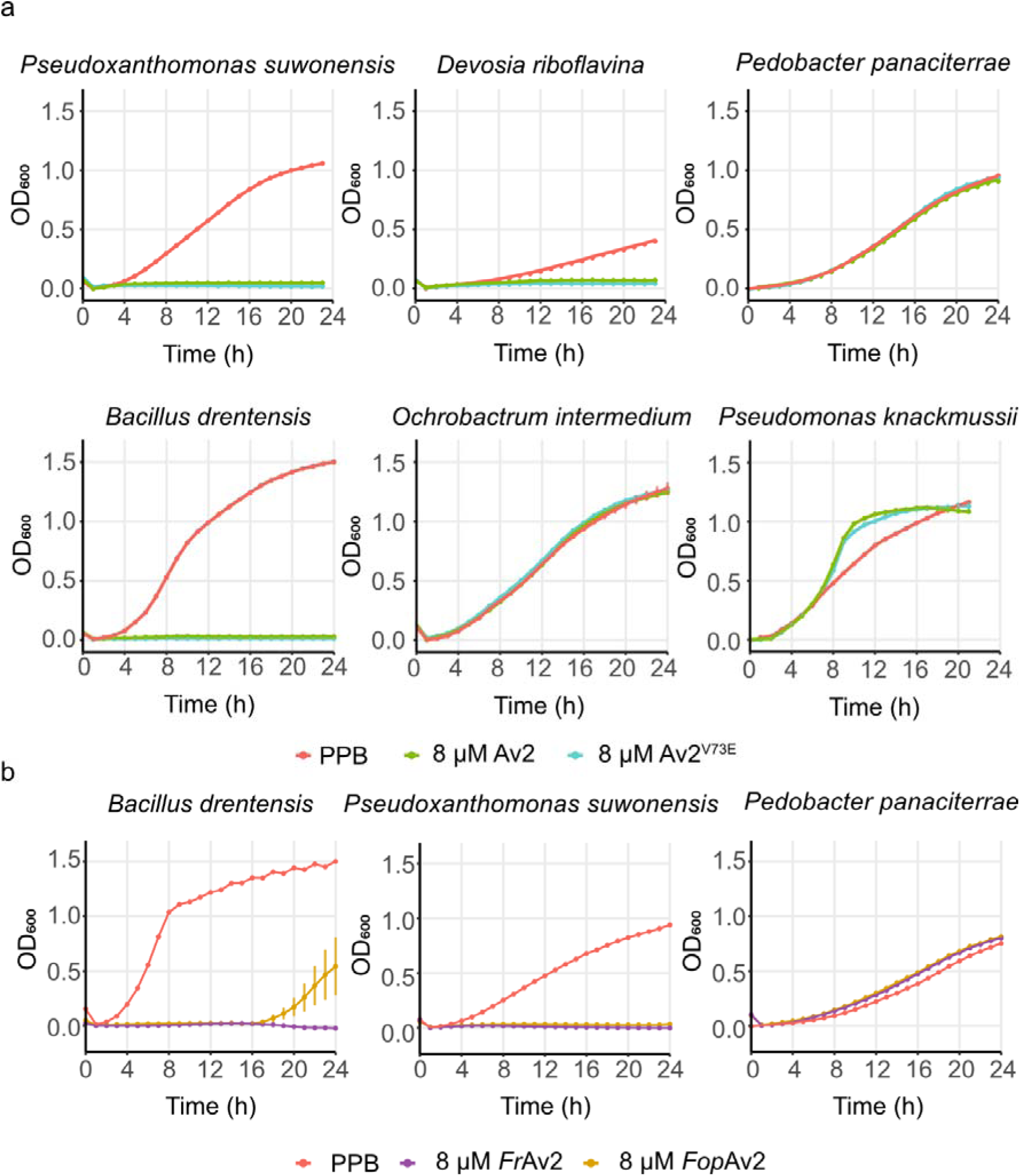
Av2 effector variants from *Verticillium dahliae* and *Fusarium* spp. display selective antibacterial activity. **(a)** The Av2 effector as well as the effector variant Av2^V73E^ selectively inhibits growth in a panel of phylogenetically diverse plant-associated bacteria *in vitro*. **(b)** Av2 homologues from *Fusarium redolans* (*Fr*Av2) and *F. oxysporum f. sp. pisi* (*Fop*Av2) display an overlapping activity spectrum with the *Verticillium dahliae* Av2 variants. Phosphate buffer (PPB) was used as control. Graphs display time-course measurements of bacterial densities in the presence or absence of effector proteins with 15 min intervals over 24 h and display the average OD_600_ of three biological replicates ± standard deviations.

### *Av2* contributes to *V. dahliae* virulence through microbiota manipulation

Next, we hypothesised that *Av2* is utilized by *V. dahliae* for microbiota manipulation during host colonization as well as during soil-colonizing stages. To investigate this hypothesis, we pursued microbiota sequencing through bacterial 16S ribosomal DNA profiling of tomato plants. To this end, tomato plants were inoculated with wild-type *V. dahliae* strain TO22 or the corresponding *Av2* deletion strain (Chavarro-Carrero et al., 2021), while water treatment was used as control. Interestingly, while tomato plants inoculated with wild-type *V. dahliae* showed severely stunted growth by ten days after inoculation when compared with mock-inoculated plants (Figure 2a), plants inoculated with the *Av2* deletion strain only showed mild symptoms of disease, and significantly less stunting occurred than in plants inoculated with the wild-type fungus. Importantly, significantly more fungal biomass was recorded in tomato plants inoculated with the wild-type fungus than in plants inoculated with the *Av2* deletion strain (Figure 2b), showing that *Av2* contributes to *V. dahliae* virulence during host colonisation.

**Fig. 2.**
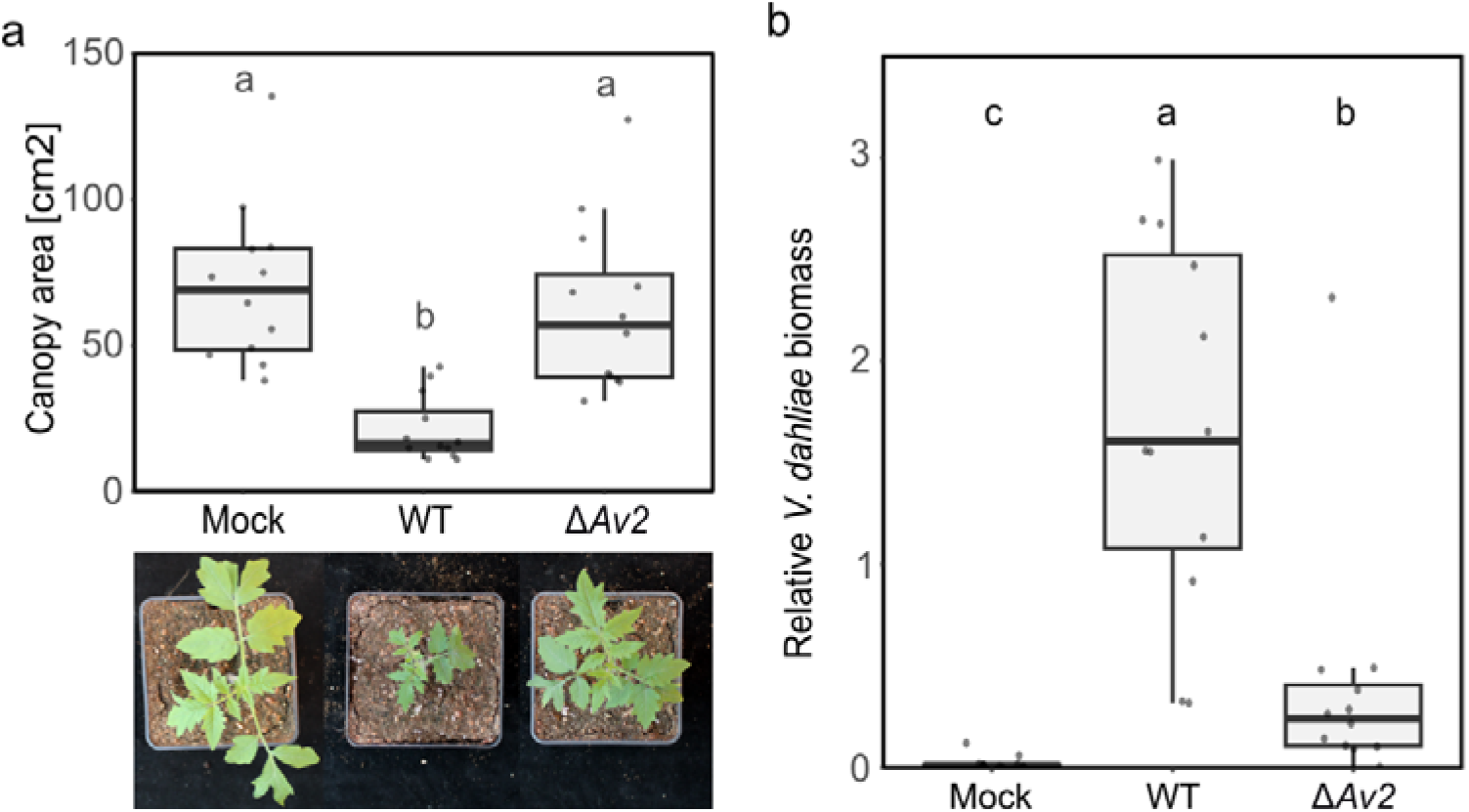
Av2 contributes to *Verticillium dahliae* virulence on tomato. **(a)** Av2 contributes to virulence of *V. dahliae* in tomato. The canopy area measurements of inoculated plants show stronger stunting upon inoculation with wild-type *V. dahliae* strain TO22 (WT) when compared with the corresponding *Av2* deletion strain (Δ*Av2*). Different letters represent significant differences (one-way ANOVA and Tukey’s post hoc test; P < 0.05). **(b)** *V. dahliae* biomass in tomato stems was quantified with real-time PCR and normalised to *Rubisco* abundance. Different letters represent significant differences (one-way ANOVA and Tukey’s post hoc test; P < 0.05).

To address the hypothesis that *Av2* contributes to virulence through microbiota manipulation, tomato plants were inoculated in a peat-based gnotobiotic system (Punt et al., 2025). If microbiota manipulation is the genuine function of the effector, *Av2* should not contribute to fungal virulence when plants are grown axenically, in the absence of microbes, while its contribution should become noticeable upon microbial reintroduction. To reintroduce microbes into sterile soil while maintaining physico-chemical properties similar to the sterilized substrate, 10% unsterilized soil was mixed with 90% sterilized soil. Importantly, plating confirmed that sterilization effectively removed the microbial population from the substrate, whereas reintroduction resulted in microbial colonization of the originally sterilized substrate (Supplementary Fig. 3). Next, tomato seedlings were inoculated with wild-type *V. dahliae* strain TO22 or the corresponding *Av2* deletion strain and cultivated in the two substrates. Importantly, at two weeks after inoculation, tomato plants inoculated with wild-type *V. dahliae* were significantly smaller than the mock-inoculated plants while plants inoculated with the *Av2* deletion strain developed similar as tomato plants grown in potting soil (Fig. 3, Fig.2a), showing that *V. dahliae* can establish infections on tomato plants also in a gnotobiotic system on sterilized substrate. As previously observed for other plant species as well, tomato plants grown axenically generally developed slower than those grown in the presence of a microbiota on recolonized substrate. Intriguingly, however, when tomato plants were grown on sterile substrate, no difference could be observed between tomato plants inoculated with wild-type *V. dahliae* or with the *Av2* deletion strain, showing that *Av2* only contributes to virulence in the presence of a microbiota. This finding suggests that microbiota manipulation is the genuine virulence function of the Av2 effector, and that the effector lacks plant virulence targets.

**Fig. 3.**
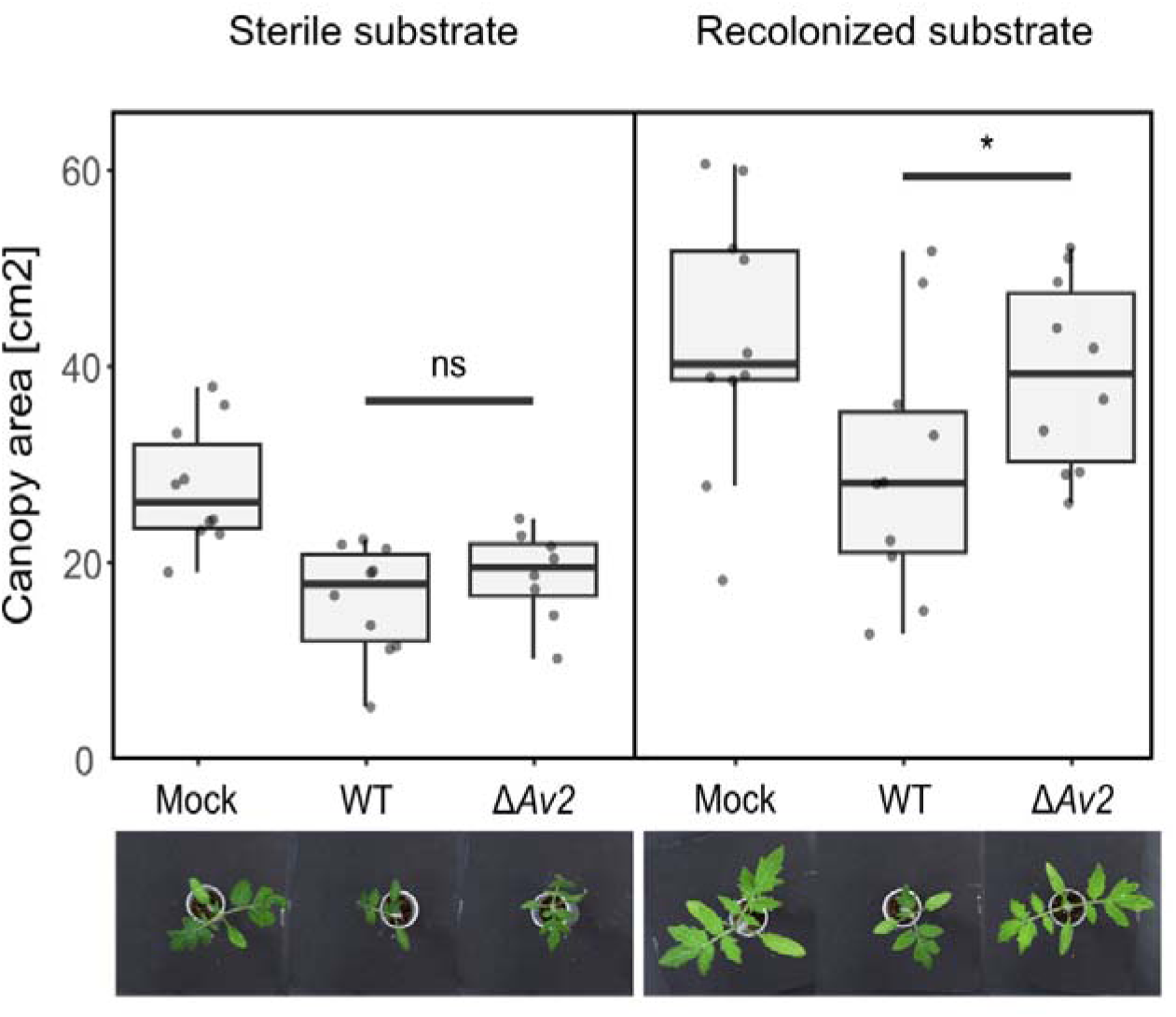
Av2 contributes to *V. dahliae* virulence on tomato plants solely in the presence of microbes. Canopy area measurements of inoculated tomato plants grown in Flowpots show stronger stunting upon inoculation with wild-type *V. dahliae* strain TO22 (WT) when compared with the corresponding *Av2* deletion strain (Δ*Av2*) in recolonized substrate but not in sterile substrate. Statistical analyses were performed for each of the substrates, and the star indicates significant differences (unpaired two-sided student’s t-test; p < 0.05). Photographs display phenotypes of representative plants for each of the treatments at 14 dpi.

### *Av2* suppresses the recruitment of Pseudomonadales

In order to perform microbiota sequencing through bacterial 16S ribosomal DNA profiling, tomato stem samples were collected at ten days post *V. dahliae* inoculation, before the onset of wilting symptoms, and the V5-V7 region of the bacterial 16S rDNA was amplified and sequenced. Subsequent analysis did not reveal major changes in microbial diversity (α- diversity) between plants inoculated with *V. dahliae* wild-type and mock-inoculated plants (Figure 4a). Interestingly, however, plants inoculated with the *V. dahliae Av2* deletion strain showed a significant reduction in microbial diversity that coincided with a strong increase in the relative abundance of Proteobacteria (Figure 4c). Principal component analysis based on weighted unifrac distance revealed differential grouping of the tomato stem endosphere microbiota for the three different treatments (PERMANOVA, P < 0.001; Figure 4b). To investigate which bacterial orders drove the separation of the samples in the principal component analysis, pairwise bacterial abundance comparisons were performed between plants inoculated with *V. dahliae* wild-type and the *Av2* deletion strain. Several bacterial orders were significantly more abundant in plants inoculated with the *Av2* deletion strain, namely Pseudomonadales, Burkholderiales, Mycobacteriales, Micromonsporales (Figure 4d). Of these bacterial orders, the Pseudomonadales displayed the largest increase in abundance (log2-fold change 1.67). Only few genera appear to drive the differential abundance of these bacterial orders. Within the Pseudomonadales only the genera *Pseudomonas* and *Acinetobacter* were significantly more abundant upon inoculation with the *Av2* deletion strain, while within the order of Burkholderiales only the genus *Massilia* showed a significant increase (Figure 4e). Especially the genus *Pseudomonas* caught our attention because of its high relative abundance in the tomato microbiota, with around 20% and 50% in plants inoculated with the wild-type *V. dahliae* and the *Av2* deletion strain, respectively. Intriguingly, while we anticipated on a reduction in *Pseudomonas* abundance in plants inoculated with wild-type *V. dahliae* when compared with mock-inoculated plants, we observed no difference in *Pseudomonas* abundance between the two treatments (Figure 4f). This significant increase of *Pseudomonas* in plants inoculated with the *Av2* deletion strain also explains the decrease in alpha diversity of this treatment (Figure 4a). Given that we only see a strong recruitment of *Pseudomonas* during the infection by the *Av2* deletion strain, we conclude that this effector is utilised by *V. dahliae* to suppress the recruitment of this bacterial genus by the host upon pathogen invasion.

**Fig. 4.**
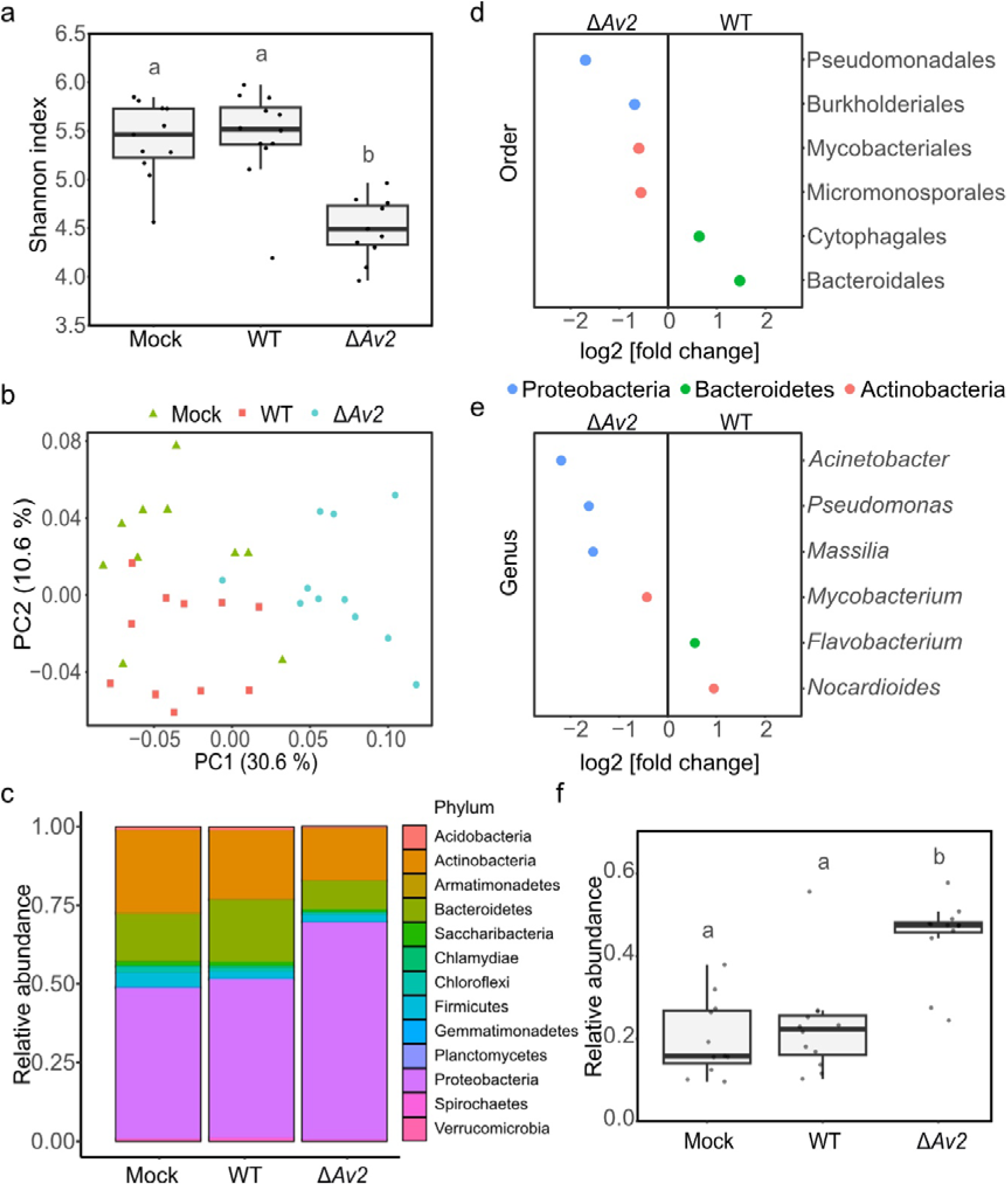
Undermining the cry for help: *Verticillium dahliae* Av2 suppresses *Pseudomonas* recruitment during host colonisation. **(a)** α-diversity of tomato endosphere microbiota ten days after inoculation as determined with 16S ribosomal DNA profiling. The α-diversity is significantly lower for microbiomes of plants inoculated with the *Av2* deletion strain when compared with the other treatments. Different letters represent significant differences (one-way ANOVA and Tukey’s post hoc test; P < 0.05). **(b)** Principal component analysis based on weighted unifrac distance reveals separation of tomato stem endosphere microbiota upon inoculation with either water (Mock), wild-type *V. dahliae* or the *Av2* deletion strain (PERMANOVA, p < 0.001). **(c)** Relative abundance of bacterial phyla show increased Proteobacteria abundance in plants inoculated with the *Av2* deletion strain. **(d)** Differentially abundant bacterial orders in the stem endosphere of tomato plants upon inoculation with either wild-type *V. dahliae* or the *Av2* deletion strain (Wald test, adjusted P < 0.05). **(e)** Differential abundance analysis of bacteria at the genus level in the tomato stems upon inoculation with either wild-type *V. dahliae* or the *Av2* deletion strain. **(f)** Relative abundance comparison of *Pseudomonas* in tomato stems upon inoculation with either water (Mock), wild-type *V. dahliae* or the *Av2* deletion strain. Different letters represent significant differences (one-way ANOVA and Tukey’s post hoc test; P < 0.05).

### *V. dahliae* utilises Av2 to inhibit antagonistic *Pseudomonas* spp

The targeted recruitment of *Pseudomonas* by tomato plants upon *V. dahliae* colonization, and the role of *Av2* in prevention of such recruitment, suggests that *Pseudomonas* acts as antagonist of the fungus. To investigate whether the interaction between *V. dahliae* and *Pseudomonas* involves direct antagonism, and to elucidate the role of *Av2* in this interaction, competition assays were performed between *V. dahliae* and *Pseudomonas* strains isolated from tomato plants (Punt et al., 2025). To this end wild-type *V. dahliae* strain TO22 and the corresponding *Av2* deletion strain were incubated with a panel of 15 *Pseudomonas* species. Interestingly, wild-type *V. dahliae* grew significantly better than the *Av2* deletion strain in presence of any of the four *Pseudomonas* species *P. laurentiana*, *P. plecoglossicida*, *P. crudilactis* or *P. vancouverensis* (Figure 5a, Supplementary Figure 3). No difference in growth between the two *V. dahliae* strains could be observed when co-cultured with the remaining *Pseudomonas* species under these conditions. The reduced growth of the *Av2* deletion strain when co-cultured with particular *Pseudomonas* species demonstrates that several *Pseudomonas* spp. are antagonists of *V. dahliae* growth and suggests that *Av2* is utilised by the fungus to counter these antagonists.

**Fig. 5.**
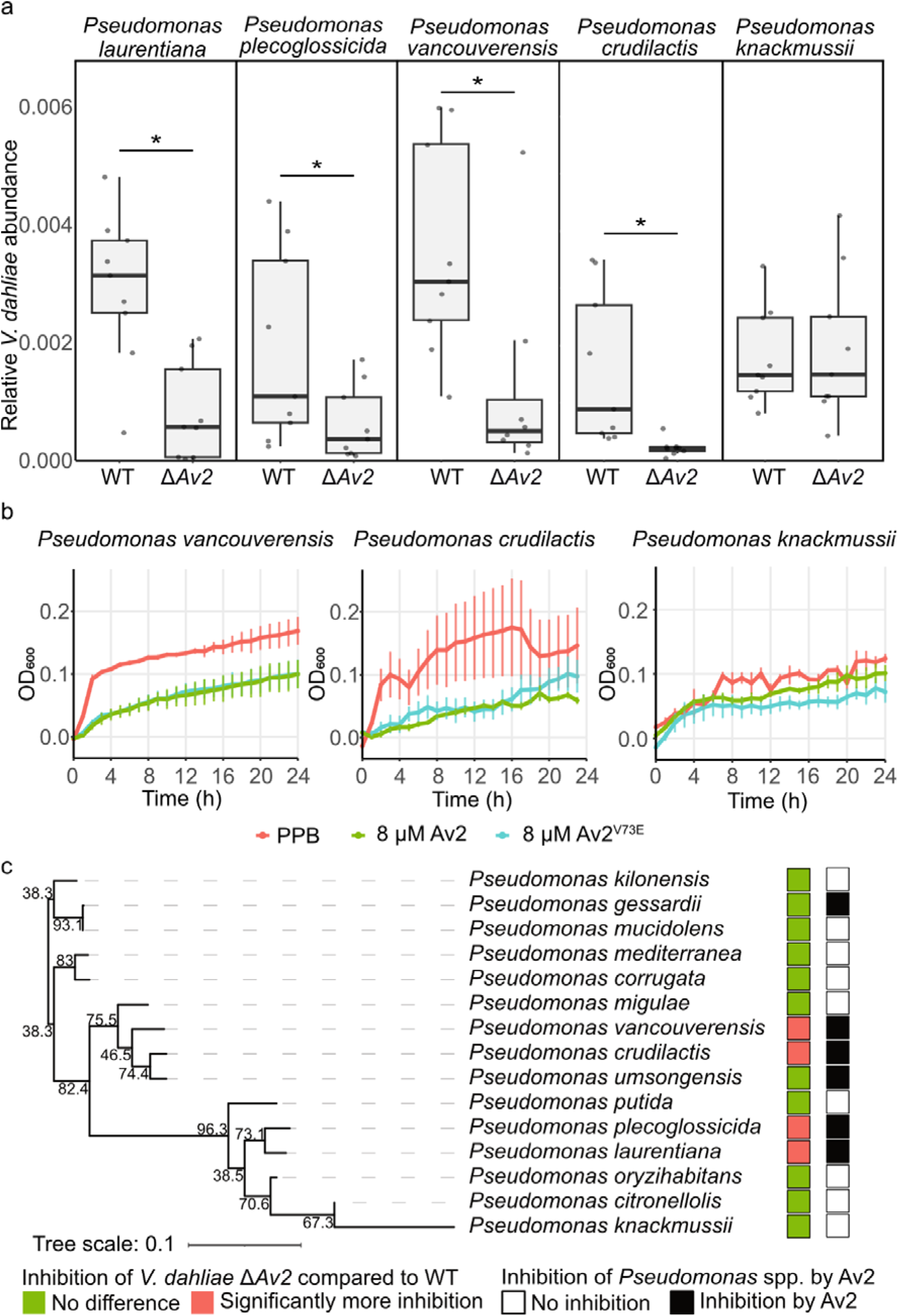
Av2 is used by *V. dahliae* for direct growth inhibition of antagonistic *Pseudomonas* spp. **(a)** Relative biomass of wild-type *V. dahliae* strain TO22 (WT) and the corresponding *VdAv2* deletion strain (Δ*Av2*) was quantified with real-time PCR after co-cultivation with a panel of Pseudomonadales in half-strength Murashige and Skoog medium for 48 h. The *V. dahliae* biomass was normalised against abundance of spike-in DNA, added during DNA extraction. **(b)** *Pseudomonas* spp. are differentially inhibited by Av2 and Av2^V73E^ *in vitro*. Phosphate buffer (PPB) was used as control. Graphs display time-course measurements with 60 min intervals over 24 h and display the average OD_600_ of three biological replicates ± standard deviations. **(c)** *Pseudomonas* spp. that display stronger antagonism towards the *Av2* deletion strain than towards wild-type *V. dahliae* do not group in a phylogenetic tree that was generated based on 2,495 orthologous genes present in all species. Inhibition of *Pseudomonas* spp. by Av2 and Av2^V73E^ *in vitro* is largely overlapping with that pattern.

To test whether Av2 inhibits the growth of antagonistic *Pseudomonas* spp., their sensitivity towards Av2 was assessed *in vitro*. Interestingly, all antagonistic *Pseudomonas* spp. that showed reduced antagonism in the presence of *Av2* were inhibited when incubated with 8 µM Av2 or Av2^V73E^ (Figure 5b). In contrast, most of the *Pseudomonas* spp. for which no difference in antagonism was recorded in the co-cultivation with *V. dahliae* were unaffected by Av2 or Av2^V73E^, suggesting that *V. dahliae* co-opted Av2 to selectively suppress antagonistic *Pseudomonas* spp (Figure 5c). To investigate the phylogenetic placement of the diverse *Pseudomonas* spp. isolates, and assess potential clustering of the species that act as *V. dahliae* antagonists and are inhibited by Av2, 2,495 orthologous genes present in all species were extracted from their genomic sequences and used to infer a phylogenetic tree. Interestingly, *Pseudomonas* spp. that showed increased antagonism towards the *Av2* deletion strain when compared with wild-type *V. dahliae* do not seem to cluster, but appear into two clades (Figure 5c). Further insight into the molecular function of Av2 could reveal whether this phylogenetic split is caused by the evolution of resistance against Av2 within the *Pseudomonas* genus or is due to physiological similarities among the inhibited antagonistic species. In conclusion, our findings suggest that *V. dahliae* exploits Av2 to supress the cry for help recruitment of beneficial *Pseudomonas* spp. during plant colonisation.

## DISCUSSION

The plant microbiota has been shown to be crucial for plant health and to act as an additional layer of defense against invading pathogens (Trivedi et al., 2020). In a phenomenon known as the “cry for help”, plants respond to pathogen invasion by dynamically altering their microbiota through modulating the composition of their root exudates to selectively recruit beneficial, disease-suppressing microorganisms and thereby mitigate disease progression (Rolfe et al., 2019). Here, we characterise the *V. dahliae* effector *Av2* as an antimicrobial effector that actively supresses the plant’s cry for help. We show that in tomato, *Av2* suppresses the recruitment of antagonistic *Pseudomonas* spp. into the rhizosphere. As a result, plants inoculated with wild-type *V. dahliae* exhibit *Pseudomonas* spp. levels comparable to mock inoculated controls, whereas infection with an *Av2*-deletion mutant leads to strong *Pseudomonas* spp. enrichment that correlates with significantly reduced fungal colonization. This activity is distinct from previously characterized antimicrobial effectors such as Ave1 and Ave1L2, which promote pathogen virulence by depleting antagonistic Sphingomonadales and Actinobacteria from the host plant microbiota (Snelders et al., 2020), or the suite of antimicrobial proteins secreted by *Albugo candida*, which collectively target core members of the *Arabidopsis thaliana* microbiota to facilitate host colonization (Gómez-Pérez et al., 2023). Our findings reveal a further sophicticated level of pathogen interference, demonstrating that pathogens can not only respond to and reshape the plant microbiome, but also sabotage microbiota-mediated host defense responses by compromising the cry for help recruitment during infection. *Pseudomonas* species are well known for their role in plant disease suppression and are frequently enriched during plant cry for help responses (Wang & Song, 2022). For example, beneficial *Pseudomonas* spp. are recruited in response to take-all disease in wheat, where they protect the host through direct antagonism against the pathogen (Raaijmakers & Weller, 1998). We observed antagonism by *P. laurentiana, P. plecoglossicida, P. crudilactis*, and *P. vancouverensis* against the *V. dahliae Av2*-deletion mutant *in vitro*, indicating that these *Pseudomonas* spp. are capable of suppressing *V. dahliae* during infection. Furthermore, the same *Pseudomonas* spp. that exhibited enhanced antagonism toward the *Av2* deletion mutant were directly inhibited by Av2. This reciprocal antagonism aligns with previous findings showing that antimicrobial effectors target beneficial bacteria that are able to antagonise the pathogen (Chavarro-Carrero et al., 2024; Snelders et al., 2020, 2023). Interestingly, *Pseudomonas* species inhibited by Av2 span two distinct phylogenetic groups, suggesting that some *Pseudomonas* species have evolved resistance to overcome suppression by Av2. This may suggest that a co-evolutionary arms race takes place between *V. dahliae* and host-associated microbiota members reminiscent of the development of antibioticum resistance. Given the abundance of microbes that produce antimicrobial molecules (Mesny & Thomma, 2024; Mullis et al., 2019), the resistance of *Pseudomonas* spp. to Av2 may be part of a broader antimicrobial resistance developed through diverse microbial interactions, with V. dahliae playing only a minor role in this process. Elucidating how particular *Pseudomonas* species have overcome Av2 sensitivity may provide valuable insight into the mode of action of the effector and selective pressures shaping pathogen–microbe interactions in the rhizosphere. Within the *V. dahliae* population, two closely related homologues of the *Av2* effector have been identified, differing by only a single amino acid (Chavarro-Carrero et al., 2021).

Since this variation does not seem to affect recognition by the V2 immune receptor (Chavarro-Carrero et al., 2021), we hypothesized that it might affect the antimicrobial activity that is exerted by the effector protein. However, our *in vitro* activity assays revealed no significant differences in antimicrobial activity between the two variants, suggesting that the amino acid substitution does not affect this function as well. *Av2* homologues have furthermore been reported in other species of the *Verticillium* genus, and in *Fusarium* (Chavarro-Carrero et al., 2021). Intriguingly, with recent evidence indicates that *V. dahliae* acquired *Av2* via horizontal gene transfer from *Fusarium* species (Sato et al., 2025). Although sequence variation exists among these homologues, our assays did not reveal any functional differences in antimicrobial activity. It is important to note, however, that only a limited panel of bacterial strains was tested, and the possibility remains that sequence variation modulates activity against untested microbial targets. The conservation of the antimicrobial function observed for Av2 is reminiscent of Ave1, which was also horizontally acquired by *V. dahliae*, in this case from plants (de Jonge et al., 2012; Snelders et al., 2020). Interestingly, plant homologues of *Ave1*, known as plant natriuretic peptides (PNP), likely exhibit similar antimicrobial activity *in vitro*, as both *Arabidopsis thaliana* PNP-A and Ave1 inhibit the growth of *Bacillus subtilis* (Snelders et al., 2020). These parallels raise the possibility that conserved antimicrobial effectors, regardless of their evolutionary origin, fulfil similar ecological roles in shaping microbial communities. Since both *Fusarium* spp. and *V. dahliae* are soil-borne fungal pathogens that infect plants via the roots and disperse within their hosts via the vasculature (Di Pietro et al., 2003; Fradin & Thomma, 2006), further investigation into the role of *Av2* in *Fusarium* spp. could help clarify whether its conserved antimicrobial activity similarly contributes to the colonization strategy shared by these pathogens.

Taken together, our findings broaden the understanding of how pathogens manipulate their hosts by revealing that antimicrobial effectors can actively suppress the pathogen-induced cry for help response. By blocking the recruitment of protective microbes, pathogens undermine a critical layer of microbiota-mediated immunity. This adds to growing evidence that the plant microbiota is a strategic battleground in host–pathogen interactions (Mesny et al., 2024). As more antimicrobial effectors are identified and characterized (Chang et al., 2021; Chavarro-Carrero et al., 2024; Gómez-Pérez et al., 2023; Kettles et al., 2018; Ökmen et al., 2023; Snelders et al., 2020, 2021, 2023) it will become increasingly clear how deeply the molecular arms race between plants and pathogens extends into the plant’s microbial sphere. Finally, given that the cry for help recruitment of beneficial microbes may ultimately lead to the establishment of disease-suppressive soils (Mesny et al., 2024), future research will have reveal whether Av2 suppresses such long-term legacy effects in the soil microbiome.

## ACKNOWLEDGEMENTS

B.P.H.J.T acknowledges funding by the Alexander von Humboldt Foundation in the framework of an Alexander von Humboldt Professorship endowed by the German Federal Ministry of Education and Research is furthermore supported by the Deutsche Forschungsgemeinschaft (DFG, German Research Foundation) under Germanýs Excellence Strategy – EXC 2048/1 – Project ID: 390686111

## AUTHOR CONTRIBUTIONS

A.K., W.P., N.C.S. and B.P.H.J.T. conceived the project. A.K., W.P., J.Z., N.C.S. and B.P.H.J.T. designed the experiments. A.K., W.P., A.D., J.Z. and N.S. performed the experiments. A.K., W.P., A.D., J.Z., N.S. and B.P.H.J.T. analyzed the data. A.K., W.P. and B.P.H.J.T. wrote the manuscript. All authors read and approved the final manuscript.

