## Supplemental Data for "Undermining the cry for help: The phytopathogenic fungus *Verticillium dahliae* secretes an antimicrobial effector protein to undermine host recruitment of antagonistic *Pseudomonas* bacteria"

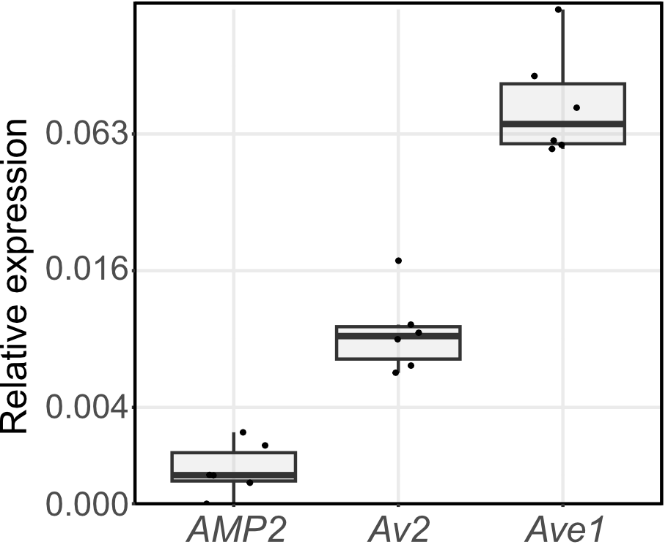

**Supplementary Fig. 1. *Verticillium dahliae* *Av2* is expressed in soil extract.** Expression of *V. dahliae* effectors after seven days of growth in soil extract when normalised to glyceraldehyde 3-phosphate dehydrogenase expression.

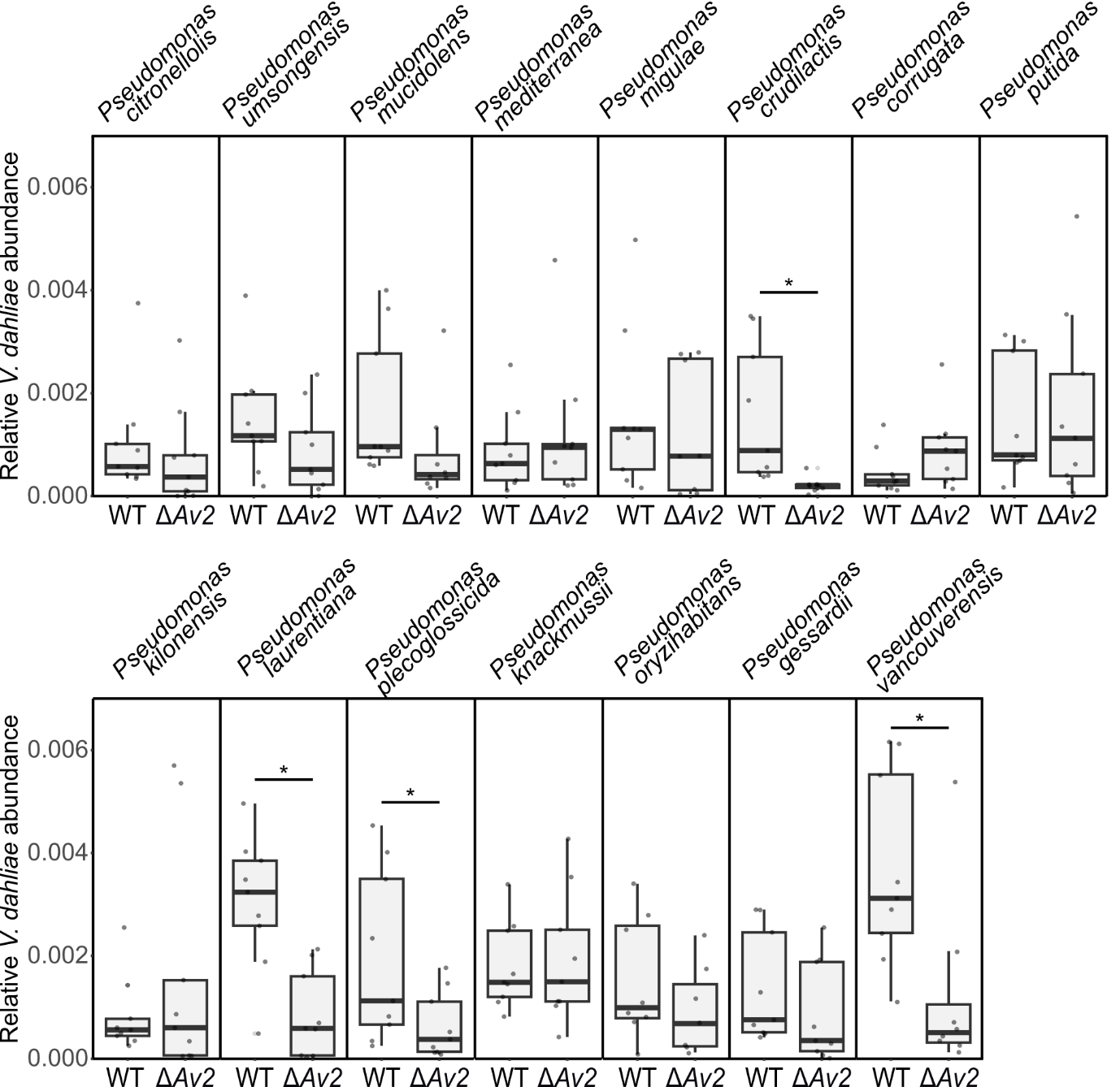

**Supplementary Fig. 2. Growth of a *Verticillium dahliae* *Av2* deletion strain is selectively impaired when co-cultured with *Pseudomonas* spp.** Relative biomass of wild-type *V. dahliae* strain TO22 (WT) and the corresponding *VdAv2* deletion strain (Δ*Av2*) was quantified with real-time PCR after co-cultivation with a panel of Pseudomonadales in half-strength Murashige and Skoog medium for 48 h*. V. dahliae* biomass was normalised against abundance of spike-in DNA added during DNA extraction.

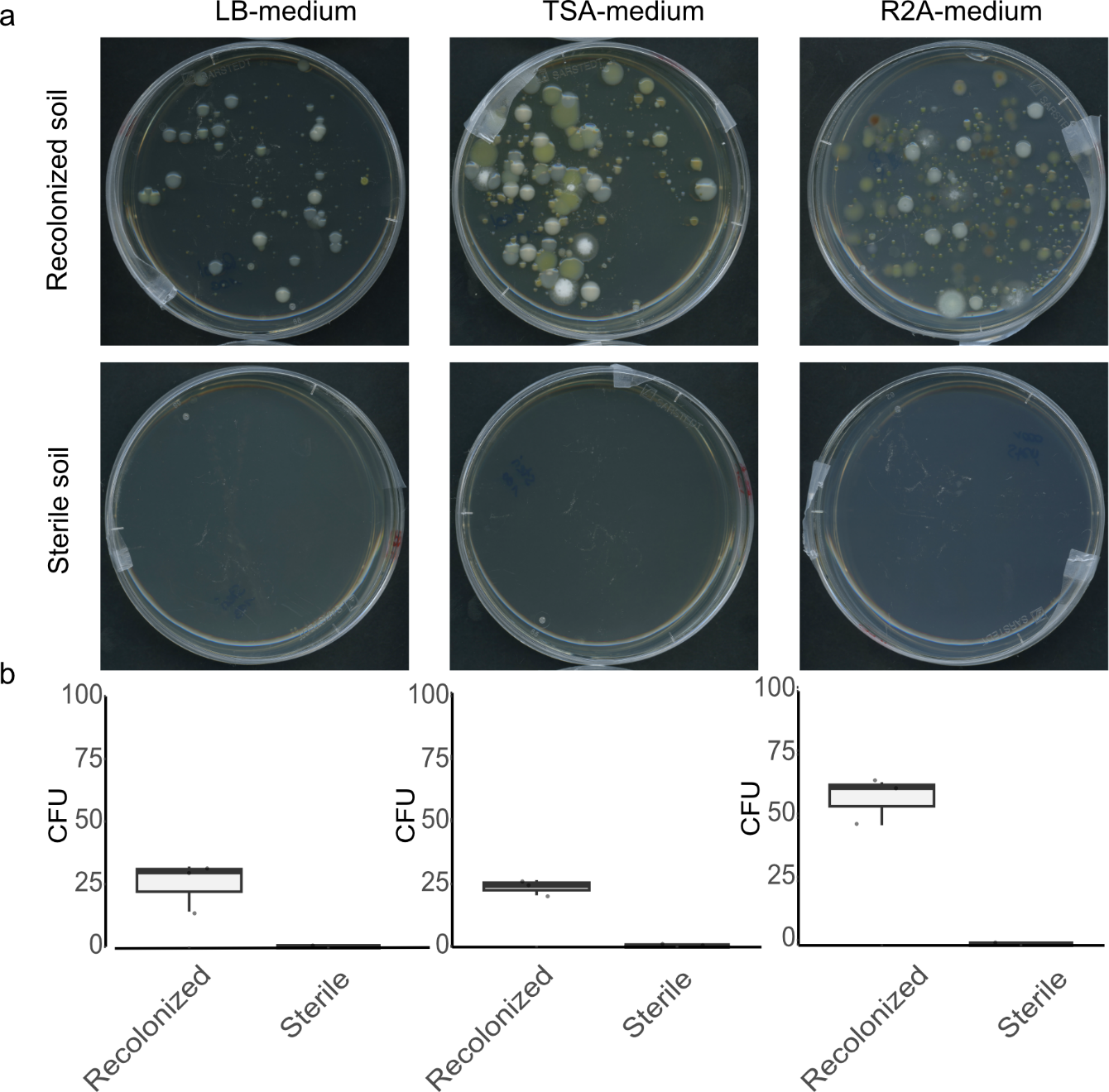

**Supplementary Fig. 3. Microbes were successfully reintroduced into sterile flowpot substrate with 10% non-autoclaved soil.** **(a)** Either recolonized or sterile Flowpot substrate were resuspended in MgCl_2_ and streaked out on three different media. There was growth on all plates containing recolonised substrate while no growth was observed on plates with sterile substrate. Photographs display agar plates after the plating of a 100x diluted substrate-MgCl_2_ suspension and a 4-day incubation in darkness at room temperature. **(b)** Boxplots displaying the number of colony-forming units (CFU) on three different growth media after plating a 100x substrate-MgCl_2_ suspension and 4 days of incubation in darkness at room temperature.
